# Higher order assembly of Sorting Nexin 16 controls tubulation and distribution of neuronal endosomes

**DOI:** 10.1101/469932

**Authors:** ShiYu Wang, Zechuan Zhao, Avital A. Rodal

## Abstract

The activities of neuronal signaling receptors depend on the maturation state of the endosomal compartments in which they reside. However, it remains unclear how the distribution of these compartments within the uniquely complex morphology of neurons is regulated, and how this distribution itself affects signaling. Here we identified mechanisms by which Sorting Nexin 16 (SNX16) controls neuronal endosomal maturation and distribution. We found that higher-order assembly of SNX16 via its coiled-coil domain drives membrane tubulation *in vitro* and endosome association in cells. In *Drosophila* motor neurons, activation of Rab5 and coiled-coil-dependent self-association of SNX16 lead to its endosomal enrichment, concomitant with depletion of SNX16-positive endosomes from the synapse, and their accumulation as Rab5- and Rab7-positive tubulated compartments at the cell body. This leads to higher levels of synaptic growth-promoting BMP receptors at the cell body, and correlates with increased synaptic growth. Our results indicate that Rab regulation of SNX16 assembly controls the endosomal distribution and signaling activities of neuronal receptors.

## Introduction

Cells detect and respond to extracellular stimuli through cell surface receptors. Ligand-bound receptors are internalized from the plasma membrane into the endosomal system, which is composed of a tubulovesicular network of polymorphic intracellular compartments. Receptors are transferred along this network in a spatially and temporally controlled manner to achieve different itineraries and specific signaling outputs (Cullen and Steinberg, 2018; Kaksonen and Roux, 2018; Rajendran et al., 2010). Neurons are highly sensitive to alterations in endosomal trafficking due to their compartmentalized morphology. The endocytic/endosomal system regulates neuronal growth and survival signaling via internalization and turnover of signaling receptors at synapses, as well as via long-distance axonal transport of receptor-containing endosomes to neuronal cell bodies (Barford et al., 2017; Deshpande and Rodal, 2016; Scott-Solomon and Kuruvilla, 2018). Regulation of these events depends on endosomal maturation, which is driven by changes in the lipid and protein composition of the endosome (Langemeyer et al., 2018).

Sorting Nexins (SNXs) are an important family of endosomal proteins, characterized by the presence of a phosphoinositide-binding Phox homology (PX) domain. SNXs participate in a variety of cellular processes, ranging from clathrin-mediated endocytosis, endosomal sorting and signaling, to lipid metabolism (Chandra and Collins, 2018; Cullen, 2008; Teasdale and Collins, 2012). These diverse roles are achieved by coordination of the PX domain with their other functional domains (Honbou et al., 2007; Lucas et al., 2016; Pylypenko et al., 2007; van Weering et al., 2012). However, the molecular and cellular mechanisms of many SNXs remain unexplored.

SNX16 has been implicated in regulating receptor traffic and signaling in both mammalian cells and in *Drosophila* (Choi et al., 2004b; Debaisieux et al., 2016; Hanson and Hong, 2003; Rodal et al., 2011). SNX16 contains a PX domain that binds specifically to PI(3)P, as well as a C-terminal coiled-coil (CC) domain, which is required for its self-association *in vivo* (Choi et al., 2004b; Hanson and Hong, 2003), and mediates dimerization in the SNX16 crystal structure (Xu et al., 2017). However, the mechanisms by which the CC domain contributes to the cellular functions of SNX16 are unknown. *Drosophila* SNX16 (dSNX16) promotes synaptic growth signaling via activated Bone Morphogenetic Protein (BMP) receptors in presynaptic terminals at the neuromuscular junction (NMJ) (Rodal et al., 2011). The F-BAR protein Nervous Wreck (Nwk) acts upstream of dSNX16 to constrain synaptic growth signaling, and this negative regulation is suppressed by mutating three highly conserved glutamates to alanines within the dSNX16 CC domain (**Fig. 1 A**; E347A, E350A, and E351A, hereafter referred to as dSNX16^3A^). Overexpression of dSNX16^3A^ in animals resulted in dramatic synaptic overgrowth and mis-localization of activated BMP receptors at the NMJ (Rodal et al., 2011), the opposite of the *dSnx16* loss-of-function phenotype, suggesting that dSNX16^3A^ is dominant active. The mechanisms by which CC-dependent activities of dSNX16^3A^ regulate neuronal endosomal traffic to promote signaling remain unclear.

**Figure 1.**
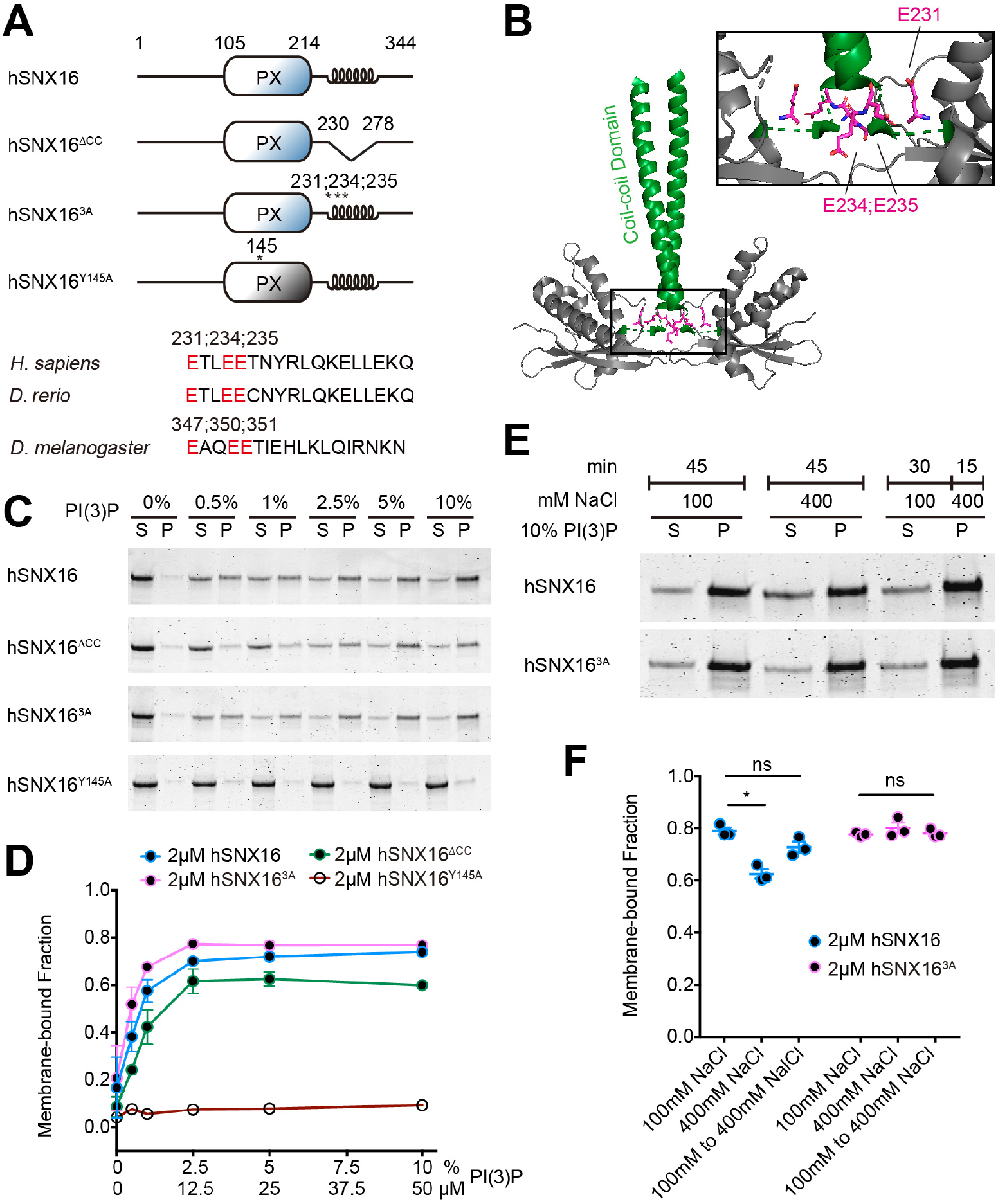
PI(3)P-dependent lipid binding of hSNX16 coiled-coil domain mutants. (A) Schematic of hSNX16 constructs. (B) Location of hSNX16 CC mutations on hSNX16 PXCC dimer (PDB# 5gw0; (Xu et al., 2017). CC domains are shown in green, and glutamates in magenta. (C-F) Liposome co-sedimentation assays. Purified hSNX16 variants were incubated with liposomes of the indicated composition and pelleted. Representative Coomassie staining of supernatant (S) and pellet (P) fractions is shown in (C) and (E). (C, D) Liposomes composed of 80% DOPC, 15% DOPE, 5% DOPS, and PI(3)P (0.5%, 1%, 2.5%, 5%, and 10%; with a corresponding decrease in PC). PI(3)P-dependent lipid binding of purified hSNX16 CC domain mutants is not significantly different from wild type hSNX16. Y145A mutation abolishes PI(3)P binding of hSNX16 as previously reported (Choi et al., 2004b). (E, F) Binding of wild type hSNX16 is more salt-sensitive than hSNX16^3A^. Liposomes (70% PC, 15% PE, 5% PS, and 10% PI(3)P) were incubated for 45 min with purified hSNX16 and the indicated NaCl concentrations before pelleting. In the last condition, hSNX16 and liposomes were incubated in 100 mM NaCl for 30 min, and then NaCl was added to a final concentration of 400 mM for 15 min before pelleting. Quantification is a result of three independent experiments, analyzed using Kruskal-Wallis test followed by Dunn’s multiple comparisons test. Data are represented as mean ± SEM. *p < 0.05.

Additional findings suggest an important role of the CC domain in SNX16 regulation. Human SNX16 (hSNX16) localizes to recycling and tubular late endosomes in addition to PI(3)P-rich early endosomes in immortalized cell lines (Brankatschk et al., 2011; Choi et al., 2004b; Hanson and Hong, 2003; Le Blanc et al., 2005; Xu et al., 2017; Zhang et al., 2013), suggesting that PI(3)P binding is not sufficient to explain its endosome association. Deleting a fragment containing CC domain results in hSNX16 loss from later endosomes and delays trafficking of internalized EGF (Hanson and Hong, 2003), highlighting the functional importance of the hSNX16 CC domain. Furthermore, structural studies reveal that PX and CC domains of SNX16 jointly contribute to entrance of PI(3)P into its binding pocket (Xu et al., 2017). Interestingly, two of the three key residues from the CC domain (E234, E235, and Y238) shaping this entrance map to E350 and E351 in dSNX16, which are mutated in dominant active mutant dSNX16^3A^ (**Fig. 1 B**; (Rodal et al., 2011; Xu et al., 2017)).

In this study, we address the role of SNX16 CC domain in regulating its molecular and cellular functions *in vitro* using purified proteins and synthetic liposomes, and *in vivo* in cultured mammalian cells and *Drosophila* motor neurons.

## Results

### hSNX16 coiled-coil variants do not strongly affect PI(3)P binding

The structure of hSNX16 suggests that its CC domain could contribute to entrance of PI(3)P into the PX domain binding pocket (Xu et al., 2017). To understand the role of the SNX16 CC domain in more detail, we purified wild type SNX16, a hSNX16 variant missing the full CC (hSNX16^∆CC^), mutant hSNX16^3A^ (E231A, E234A and E235A), as well as a PX domain PI(3)P-binding pocket mutant hSNX16^Y145A^ (Choi et al., 2004b) as a negative control (**Fig. 1, A and B**). We then tested PI(3)P binding of different CC variants in liposome co-sedimentation assays (**Fig. 1 C and D**). Wild type hSNX16 co-sedimented more effectively with increasing PI(3)P concentration, and hSNX16^Y145A^ abolishes PI(3)P binding, as previously shown (Choi et al., 2004b). By contrast, PI(3)P binding sensitivities of hSNX16^∆CC^ and hSNX16^3A^ were not significantly different from hSNX16 (**Fig. 1 D**), suggesting that the CC domain does not strongly modulate the lipid-binding affinity of SNX16. We next tested if electrostatic interactions mediate the binding of hSNX16 and our CC variants to the membrane by conducting liposome co-sedimentation assays in the presence of 400 mM NaCl. hSNX16 binding to 10% PI(3)P liposomes was reduced in the presence of NaCl. However, once bound to liposomes, hSNX16 was resistant to 400 mM NaCl, suggesting that the initial binding step is sensitive to electrostatic interactions, followed by more hydrophobically-driven effects. Interestingly, though hSNX16^3A^ exhibited similar lipid binding to hSNX16 at 10% PI(3)P, hSNX16^3A^ binding was not sensitive to 400 mM NaCl (**Fig. 1, E and F**), suggesting that the CC domain may be involved in these hydrophobic interactions. These results raise the possibility that the CC domain of hSNX16 functions in parallel to the canonical PI(3)P-binding PX domain to mediate membrane association.

### hSNX16 oligomerizes into higher-order assemblies via its coiled-coil domain

Since the hSNX16 CC domain mediates its homo-oligomerization (Choi et al., 2004b; Hanson and Hong, 2003), we hypothesized that altered salt sensitivity of hSNX16^3A^ may be due to changes in self-association. To test this hypothesis, we incubated hSNX16 CC variants with 1% PI(3)P liposomes and increasing concentrations of the 8-atom-long lysine crosslinker BS^3^, and conducted co-sedimentation assays (**Fig. 2, A and B**). In the liposome-containing pellet, hSNX16 exhibited crosslinked species at the molecular weight of dimers (~100 kDa, white triangle) as well as unexpected higher-order species (>200 kDa, red triangle). hSNX16^∆CC^ produced minimal higher-order species, though we did observe dimers, consistent with previous findings that the PX domain contributes to self-association (Choi et al., 2004b). By contrast, hSNX16^3A^ exhibited more abundant crosslinking than wild type hSNX16 into higher-order species (**Fig. 2 A**), though total co-sedimentation with PI(3)P was similar to hSNX16 and hSNX16^∆CC^ (**Fig. S1, A and B**). In the supernatant fraction, we also observed higher-order assemblies of hSNX16 and hSNX16^3A^, and only dimer species of hSNX16^∆CC^ (**Fig. 2 B**). Interestingly, hSNX16^3A^ supernatants lacked the dimer species and only showed higher order assemblies. We then asked if these oligomers required the presence of membranes by conducting crosslinking assays in buffer alone. In contrast to liposome-containing crosslinking experiments, in the absence of liposomes hSNX16 and hSNX16^3A^ showed limited, comparable dimer and higher order assembly formation (**Fig. S1 C**), suggesting that oligomerization of hSNX16^3A^ is potentiated by membranes. Taken together, we conclude that SNX16 can oligomerize into higher order assemblies via its CC domain, and that hSNX16^3A^, a mutant in the CC domain, promotes formation of these assemblies.

**Figure 2.**
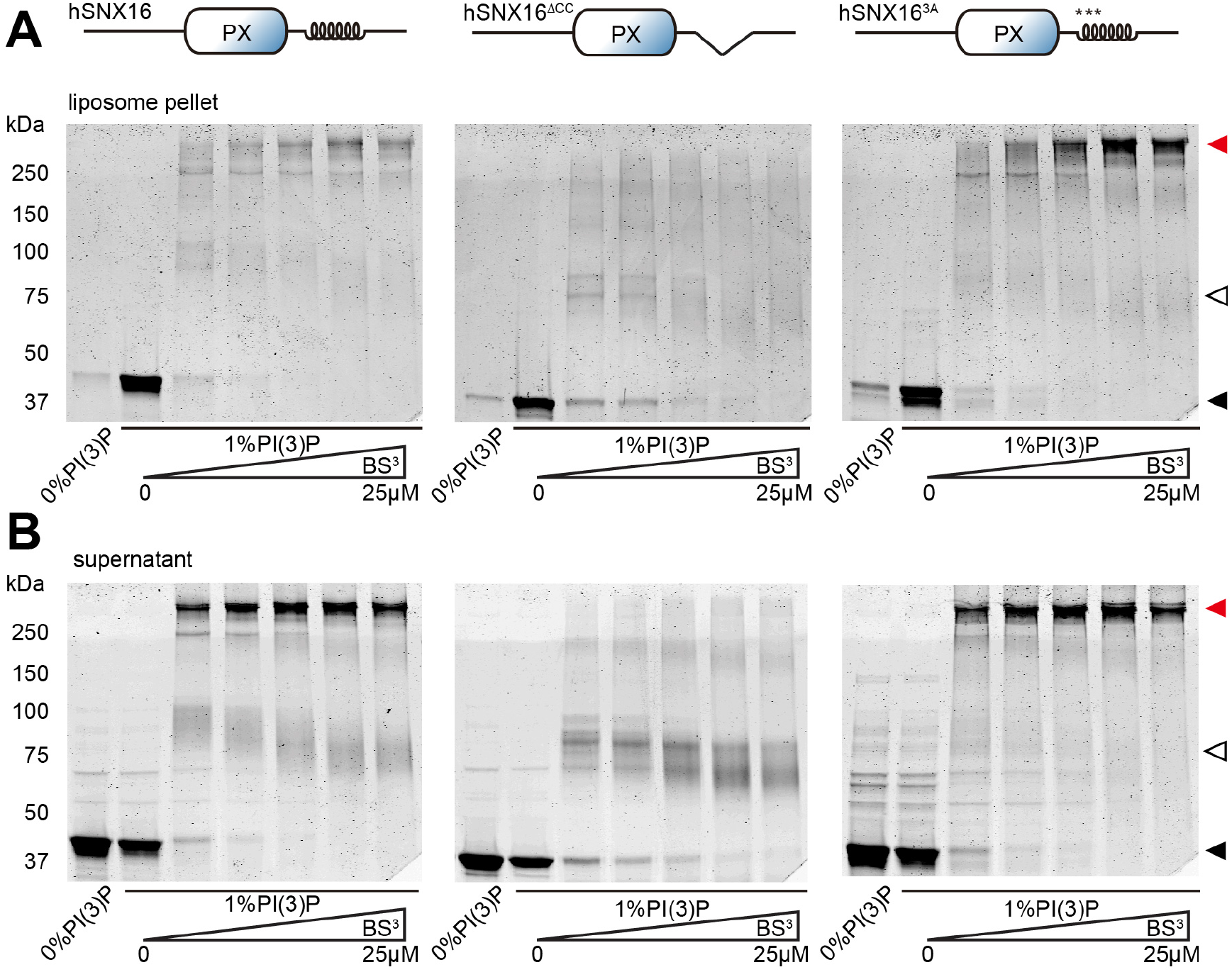
hSNX16^∆CC^ reduces and hSNX16^3A^ promotes self-association compared to hSNX16^WT^ on PI3P liposomes. (A-B) 5 µM purified hSNX16, hSNX16^3A^, or hSNX16^∆CC^ were incubated with 1% (5 µM) PI(3)P liposomes followed by BS^3^ crosslinking, and then tested for liposome co-sedimentation. (A) Coomassie-stained gels of crosslinked high-molecular-weight assemblies in protein-liposome pellets with increasing BS^3^ concentration (125 nM, 2.5 μM, 5 μM, 12.5 μM, and 25 μM). (B) Coomassie-stained gels of unbound proteins in the lipid co-sedimentation assay from same experiments as in (A). Black, white, and red triangles point to monomers, dimers, and higher-order assemblies, respectively.

To further explore the formation of hSNX16 higher-order assemblies, we conducted fluorescence recovery after photobleaching (FRAP) experiments on giant unilamellar vesicles (GUVs) coated with SNAP^488^-labeled hSNX16 variants (**Fig. 3 A**). All hSNX16 variants coated GUVs uniformly and did not exhibit localized clustering. At 100 nM, hSNX16^3A^ failed to recover after photobleaching a region of the GUV, while hSNX16 recovered rapidly (**Fig. 3, B and C**). At 500 nM, hSNX16 and hSNX16^3A^ both failed to recover after photobleaching, while hSNX16^∆CC^ remained mobile, suggesting that hSNX16 higher-order assemblies form a stable scaffold on GUV via the CC domain, and that the hSNX16^3A^ mutant promotes these stable assemblies, consistent with our crosslinking results (**Fig. 3 C and S2 A**). In addition, labeled phosphatidyl-ethanolamine (PE) in the photobleached region recovered rapidly for all hSNX16 CC variants, indicating that this component of the membrane remained fluid even when the protein assembly did not (**Fig. 3 D**). Considering both the crosslinking and the GUV FRAP results, we conclude that hSNX16 can oligomerize into higher-order structures (beyond dimers) via its CC domain, and that the hSNX16^3A^ mutant and membrane binding together increase formation of these oligomers.

**Figure 3.**
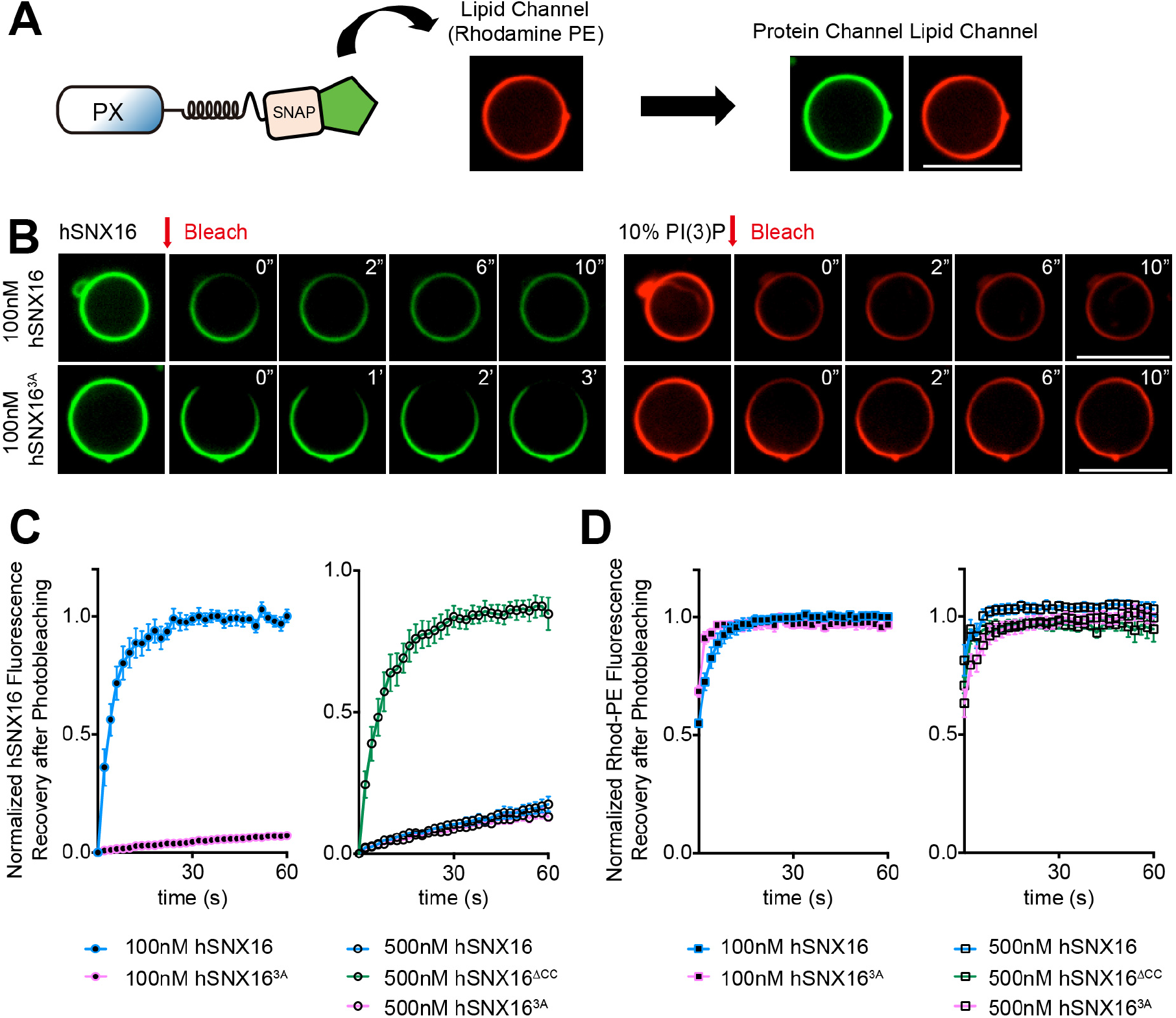
hSNX16 oligomerizes into higher-order assemblies on membranes via its coiled-coil domain. (A) Schematic of GUV assay. (BD) hSNX16^3A^ fails to recover after photobleaching at lower protein concentrations compared to wild type hSNX16, while hSNX16^∆CC^ retains high mobility at all concentrations measured. (B) Representative time-lapse images of 100 nM hSNX16 and hSNX16^3A^ before and after photobleaching. (C-D) Quantification of protein and lipid fluorescence of GUVs bound by hSNX16 variants at 100 nM and 500 nM. Protein and lipid fluorescence were normalized to a non-bleached region on the same GUV to correct for photobleaching. Protein fluorescence was then further normalized by subtracting from all timepoints the intensity at t=0, and normalizing prebleach intensity to 1. Quantification is a result of at least five independent GUVs. Data are represented as mean ± SEM. Scale bar, 10 μm.

### hSNX16 higher order assemblies promote membrane tubulation

When conducting the FRAP experiments with 500 nM hSNX16 variants, we consistently observed tubule formation on hSNX16 and hSNX16^3A^, but not hSNX16^∆CC^ coated GUVs (**Fig. S2 A**). This was unexpected since SNX16 does not contain a BAR-domain scaffold like other membrane-tubulating sorting nexins (van Weering et al., 2012), and instead forms a scissor-shaped homodimer (Xu et al., 2017). To investigate this novel membrane tubulation activity of hSNX16, we incubated hSNX16 CC variants with GUVs composed of various PI(3)P compositions, and quantified GUV morphology (**Fig. 4, A and B**). We did not observe tubules on 0% PI(3)P GUVs incubated with hSNX16, or on GUVs without any protein. By contrast, hSNX16 generated tubules on 10% PI(3)P GUVs at 100 nM, and the number of tubulated GUVs increased at 500 nM hSNX16 (**Fig. 4, A and B**). Strikingly, hSNX16^3A^ generated tubules at lower PI(3)P and protein concentrations than hSNX16, and hSNX16^∆CC^ failed to generate tubules under any tested conditions (**Fig. 4, A and B**). Thus, the tubulation activity of hSNX16 and our CC variants correlated with their capability to form higher-order assemblies in cross-linking and FRAP assays (**Figs. 2 and 3**).

**Figure 4.**
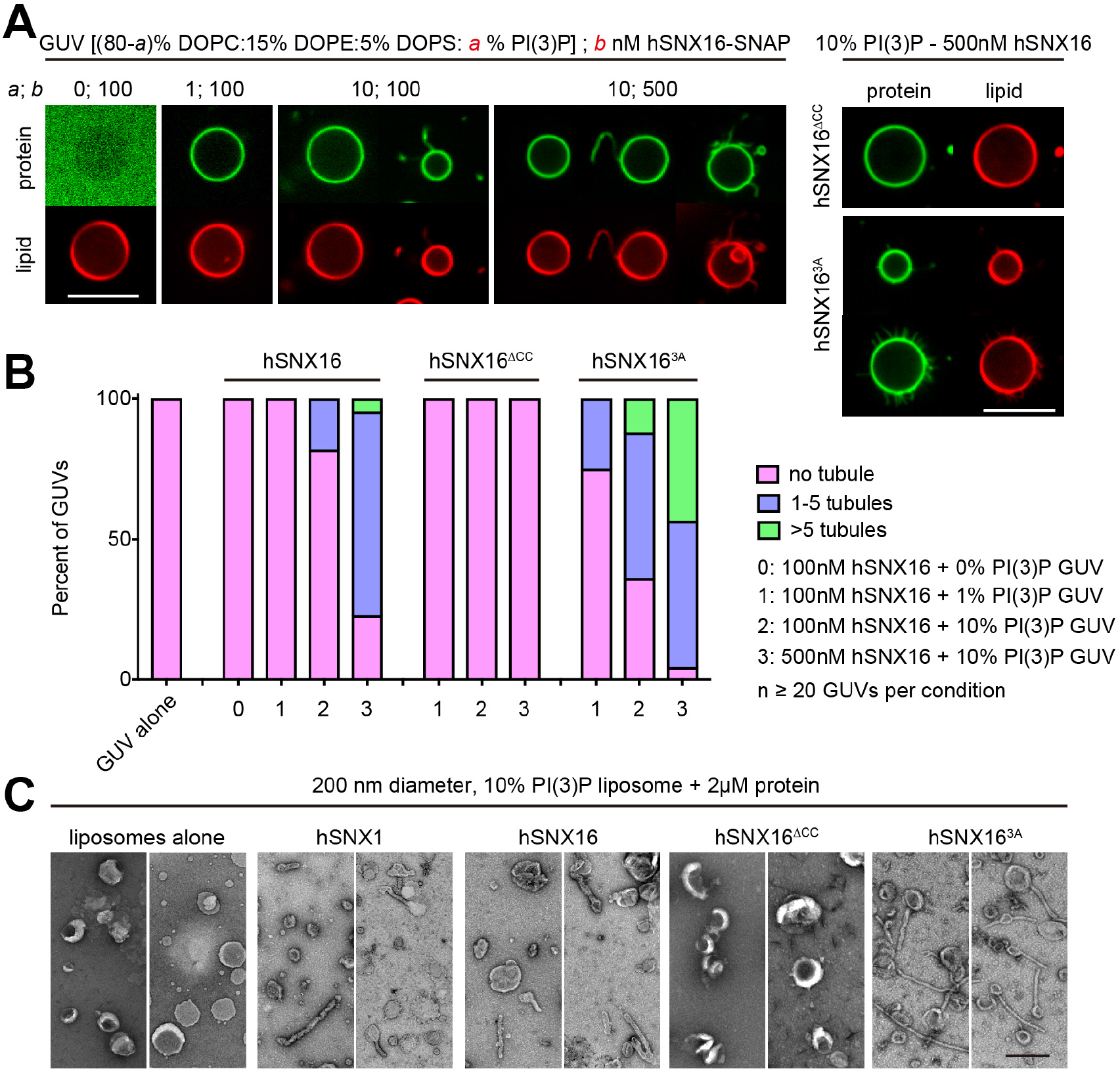
hSNX16 generates membrane tubules via its coiled-coil region. (A) Representative single confocal slices of Rhod-PE labeled GUVs (0%, 1% (30 nM) or 10% (300 nM) PI(3)P) incubated with 100 nM or 500 nM hSNX16 CC variants. Scale bar, 10 μm. (B) Percentage of tubulated vesicles at indicated PI(3)P and hSNX16 concentrations. Quantification is from at least 20 GUVs per condition. (C) Negative stain EM of liposomes incubated with buffer, hSNX1, or hSNX16 CC variants (2 μM final protein concentration). Two representative fields are shown for each condition. Scale bar, 400 nm.

Because GUVs are cell-size vesicles that are essentially flat at the scale of individual proteins, we next tested whether hSNX16 promotes membrane tubulation on smaller, endosome-size vesicles using negative stain electron microscopy (EM). After incubation with 200 nm liposomes, hSNX16 and hSNX16^3A^ exhibited comparable tubulation to the BAR domain-containing sorting nexin 1 (SNX1; **Fig. 4 C**). Thus, hSNX16 generates membrane tubules via its CC domain, and this membrane tubulation activity correlates with higher-order assembly. Taken together, our *in vitro* data indicate that the CC domain is required for hSNX16 higher-order assembly and membrane deforming activities, and that the hyper-oligomerized mutant hSNX16^3A^ promotes these activities.

### hSNX16 coiled-coil variants control endosome association in vivo

To investigate the importance of hSNX16 oligomerization *in vivo*, we next tested the subcellular localization of hSNX16 CC variants. hSNX16 exhibited both punctate and cytoplasmic localization, and some puncta were elongated or tubulated, as previously reported (**Fig. 5**) (Brankatschk et al., 2011; Choi et al., 2004b; Xu et al., 2017). One previous study found that deletion of a hSNX16 fragment longer than but including the CC (aa214-aa295) resulted in its loss from late endosomes while preserving early endosome localization in A431 cells (Hanson and Hong, 2003). We further found that a more precise deletion of the CC domain (aa230-aa278) resulted in partial loss of SNX16 from punctate structures and net redistribution to the cytoplasm in cultured rat hippocampal neurons (**Fig. 5 A**). By contrast, hSNX16^3A^ exhibited enhanced punctate localization and was depleted from the cytosol. To understand whether this subcellular localization of hSNX16 CC variants is unique to neurons, we tested multiple mammalian cell lines and observed similar results, suggesting a general role for CC domain-promoted oligomerization in hSNX16 endosomal localization (**Fig. 5 B**). We further quantified the change in hSNX16^3A^ subcellular localization and found that it was significantly different from hSNX16, reflected both in the increase of CoV at the neuronal cell body and a larger fraction of high intensity pixels in HeLa cells (**Fig. 5, C and D**). Thus, we conclude that hSNX16 CC domain is required for its proper endosomal localization *in vivo*, and that the hSNX16^3A^ mutant, which exhibits enhanced oligomerization *in vitro, also* promotes SNX16 endosome localization.

**Figure 5.**
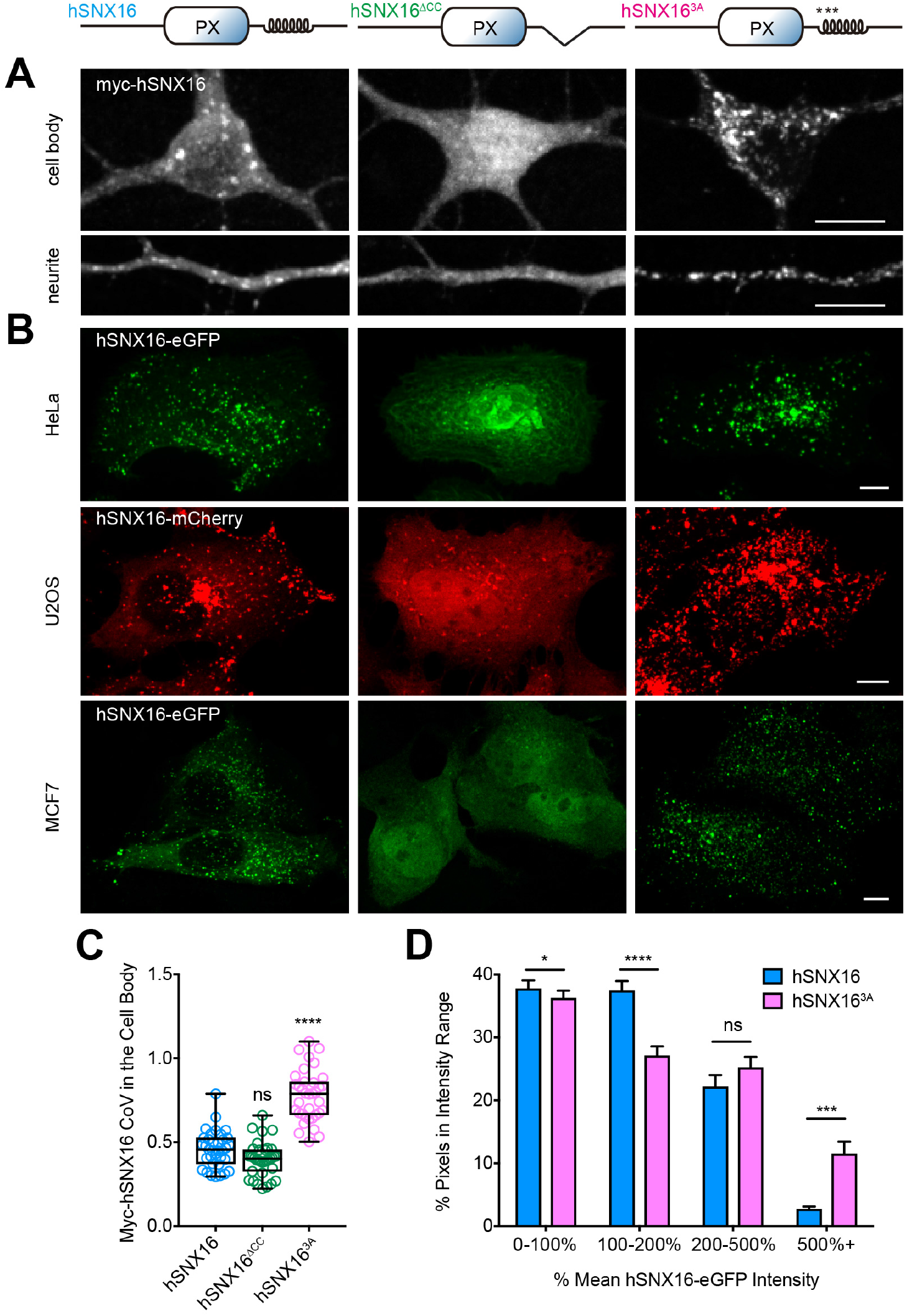
hSNX16 coiled-coil mutants exhibit altered subcellular localization *in vivo*. hSNX16^∆CC^ reduces and hSNX16^3A^ enhances hSNX16 punctate localization in mammalian cells. (A) Representative images of cell body and neurites from immunostained DIV 7 rat hippocampal neurons transiently expressing the indicated myc-hSNX16 variants. (B) Representative images of the indicated cell lines expressing indicated hSNX16 CC variants and fixed 24 hours after transfection. (C) Coefficient of variance (CoV) of myc-hSNX16 CC mutants in hippocampal cell bodies. Quantification is from at least 35 neurons per condition from three independent coverslips, analyzed using one-way ANOVA followed by Tukey’s Test. (D) Histograms depict fraction of pixels at indicated intensities for hSNX16 and hSNX16^3A^ in HeLa cells. Quantification is from at least 14 cells per condition, analyzed using Mann-Whitney test within each bin. All images show 2D maximum intensity projections of confocal stacks. Data are shown as box-and-whisker plots with all data points superimposed in (C), and as mean + SEM in (D). *p < 0.05, ***p < 0.001, ****p < 0.0001. Scale bar, 10 μm.

### dSNX16 coiled-coil variants control localization and tubulation of compartments in Drosophila motor neurons

Previously, we reported that null mutants of *dSnx16* cause reduced synaptic growth at the *Drosophila* NMJ, and that dSNX16^3A^ acts as a dominant active mutant and exhibits a synaptic overgrowth phenotype (Rodal et al., 2011). We therefore examined how the SNX16 CC-mediated oligomerization behavior that we observed in the mammalian system could contribute to its functional role at the *Drosophila* NMJ. We first tested whether the *in vivo* change in subcellular localization that we observed in mammalian cells is conserved. We generated transgenic fly lines with SNAP-tagged dSNX16 CC variants, each inserted into the same genetic locus, and used the binary GAL4/UAS system to examine their localization upon expression in *Drosophila* larval motor neurons. This allowed us to examine localization within different neural structures, including cell bodies and dendrites in the ventral ganglion, axons bundled in segmental nerves, and NMJ axon terminals embedded at the surface of the muscle (See **Fig. S3 A** for schematic). dSNX16 primarily localized to punctate structures at the NMJ (**Fig. 6 A**), which, as we previously described, correspond to endosomes (Rodal et al., 2011). Further, we found that similar to hSNX16 localization in mammalian cells, dSNX16 exhibited both cytoplasmic and punctate localization in axons proximal to the ventral ganglion, as well as in the cell bodies. Deletion of the dSNX16 CC resulted in increased cytoplasmic localization, while dSNX16^3A^ exhibited enhanced punctate localization, similar to our observations for hSNX16 (**Fig. 6, A and C**), indicating that the function of the CC domain in SNX16 localization is conserved. Furthermore, via structure illumination microscopy (SIM) of neuronal cell bodies, we found that dSNX16 localized to tubulated structures (**Fig. 6 B**). This tubule localization was CC domain dependent, and an increased number of tubulated compartments were observed in the cell body of dSNX16^3A-^expressing animals, correlating with the tubulation activity of hSNX16 CC variants *in vitro*. We next measured the intensity of dSNX16 CC variants at the NMJ, proximal axon, and cell body, and found that dSNX16^3A^ was specifically enriched at the cell body, while dSNX16^∆CC^ exhibited overall decreased intensity (**Fig. 6 D**). The enhanced punctate localization and cell body enrichment of dSNX16^3A^ was also observed for randomly integrated UAS-dSNX16-GFP P-element lines with different expression levels (**Fig. S3, B-D**; (Rodal et al., 2011)). These results suggest that in *Drosophila* motor neurons, where SNX16 positively regulates synaptic growth signaling, dSNX16 CC variants control the abundance, distribution, and structure of endosomes.

**Figure 6.**
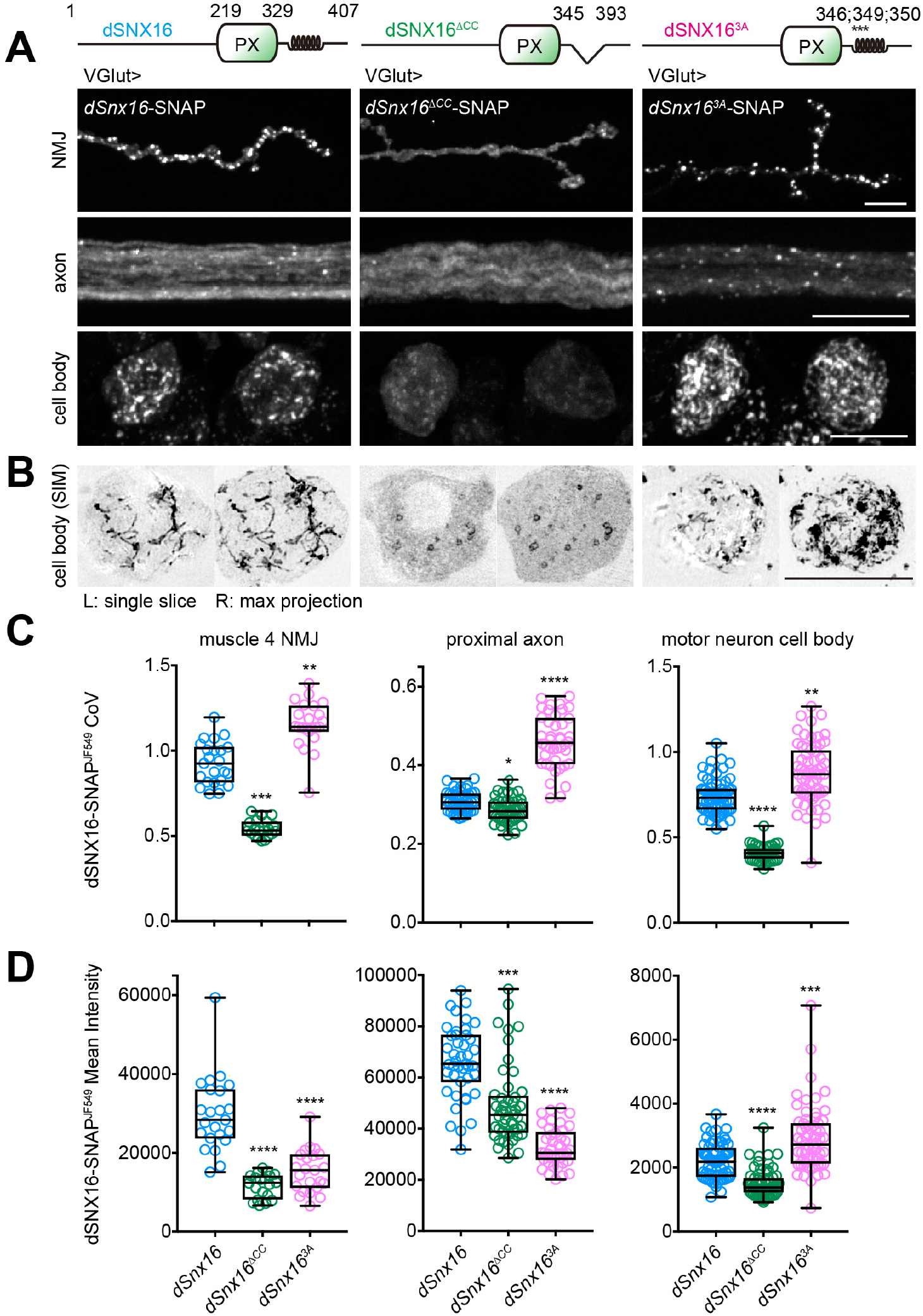
dSNX16 coiled-coil variants alter endosome structure, localization and distribution in larval motor neurons. (A-B) Representative images of animals expressing indicated UAS-dSnx16-SNAP variants driven by VGlut-GAL4. Shown are muscle 4 NMJ, proximal axons (within 100 µm of the ventral ganglion), and MNISN-I cell bodies. dSNX16^∆CC^ reduces and dSNX16^3A^ enhances dSNX16 punctate localization. dSNX16^3A^ levels are increased at the cell body and reduced at the NMJ. (B) dSNX16 localizes tubular structures at the cell body revealed by structured illumination microscopy (SIM). dSNX16^∆CC^ reduces and dSNX16^3A^ increases the quantity of tubulated SNX16 compartments. (C-D) Coefficient of variance (CoV) and mean intensity quantification of dSNX16-SNAP^JF549^. Quantification is from at least 20 NMJs, 42 axons, or 65 cell bodies, analyzed using Kruskal-Wallis test followed by Dunn’s multiple comparisons test. All images show 2D maximum intensity projections of confocal stacks unless noted otherwise. Data are shown as boxand-whisker plots with all data points superimposed. *p < 0.05, **p < 0.01, ***p < 0.001, ****p < 0.0001. Scale bar, 10 μm.

### dSNX16^3A^ enriches the BMP receptor Thickveins at the cell body

dSNX16 colocalizes with the BMP signaling pathway receptor Thickveins (Tkv) at the NMJ (Rodal et al., 2011). To understand the relevance of enhanced punctate localization and cell body enrichment of dSNX16^3A^, we examined the colocalization between GFP-tagged dSNX16 and dSNX16^3A^, and mCherry-tagged Tkv throughout the entire neuron (**Fig. 7 A**), using two different dSNX16^3A^-GFP lines to control for the effects of expression levels. Colocalization of dSNX16 with Tkv was increased in both high and low-expressing dSNX16^3A^-GFP lines, concomitant with increased Tkv intensity at the cell body (**Fig. 7, C and D**). SIM further revealed that Tkv localized within SNX16-coated endosomes, but was strikingly absent from SNX16-decorated tubules (**Fig. 7 B**, arrows). The stronger dSNX16^3A^ line also exhibited enhanced colocalization with Tkv at the NMJ and the proximal axon with overall elevated Tkv intensity, while the weaker dSNX16^3A^ line did not change Tkv localization or intensity in these regions of the neuron (**Fig. 7, C and D**), suggesting that BMP signaling components at the cell body are more sensitive to the effects of dSNX16^3A^. Taken together, these results suggest that dSNX16 CC-mediated endosome association is important for the levels and distribution of the BMP receptor Tkv in motor neurons.

**Figure 7.**
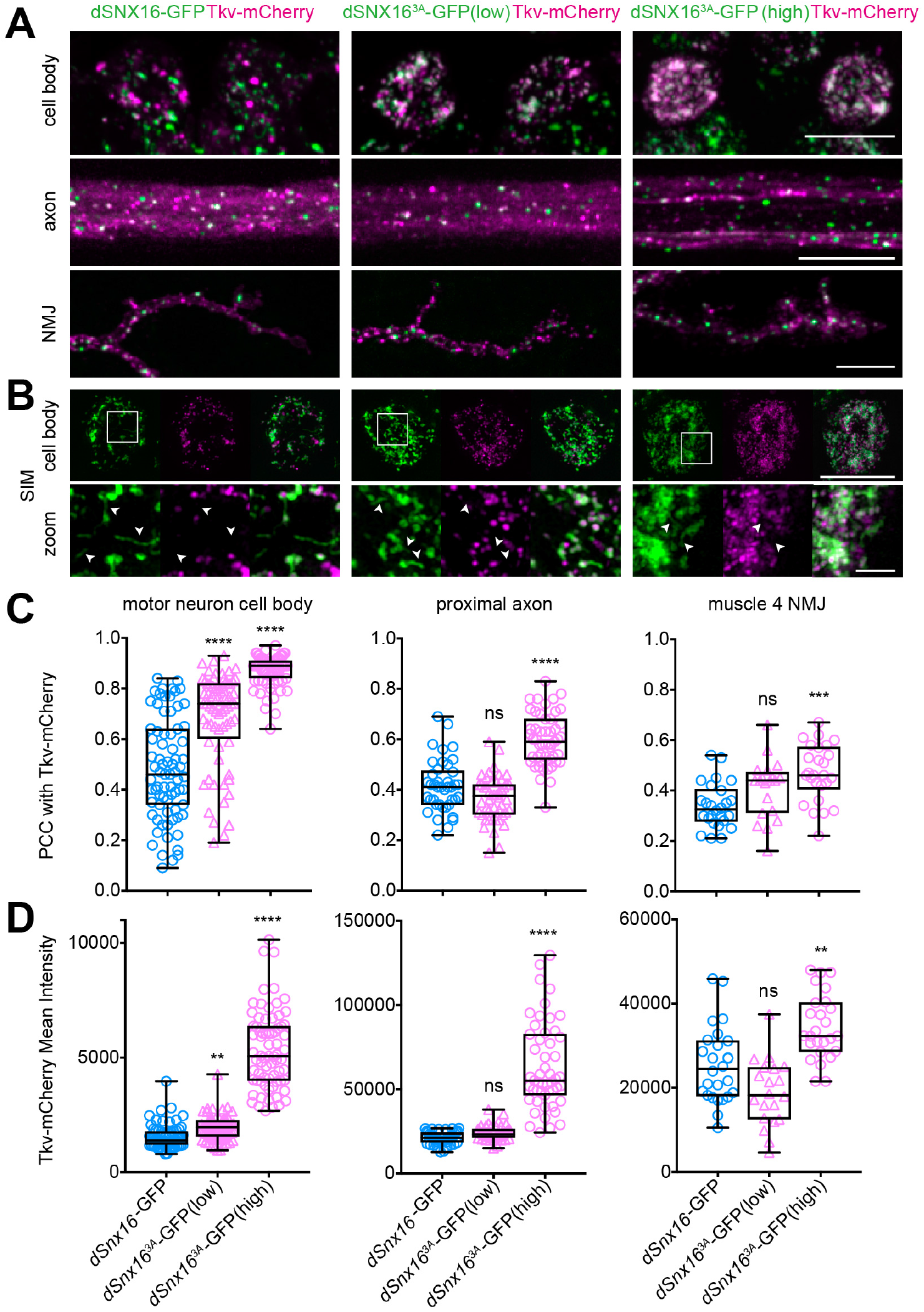
dSNX16^3A^ increases colocalization with and causes accumulation of the BMP receptor Tkv at the cell body. (A-B) Representative images of animals expressing VGlut-driven Tkv-mCherry and the indicated dSNX16-GFP transgenes at muscle 4 NMJ, proximal axon, and in the cell body. dSNX16^3A^-GFP (low) and dSNX16^3A^-GFP (high) lines correspond with dSNX16^3A-^GFPIIA and dSNX16^3A^-GFPIIIF lines in Fig. S3. Images were acquired and quantified with identical settings but are shown with contrast adjusted. White arrows in (B) point to tubular dSNX16 compartments that do not contain Tkv. (C) Pearson correlations between TkvmCherry and dSNX16-GFP. (D) Mean intensity quantification of Tkv-mCherry. Quantification is from at least 20 NMJs, 48 axons, or 72 cell bodies, analyzed using Kruskal-Wallis tests followed by Dunn’s multiple comparisons test. All images show 2D maximum intensity projections of confocal stacks. Data are shown as boxand-whisker plots with all data points superimposed. **p < 0.01, ***p < 0.001, ****p < 0.0001. Scale bar, 10 μm, or in the zoomed-in view of (B), 2 μm.

### dSNX16 associates with different endosomal compartments at the NMJ compared to the cell body

Our results indicate that the dSNX16 CC may have distinct effects on endosomes in different regions of neurons. To further understand the nature of dSNX16 puncta in these regions, we examined the co-localization dSNX16 CC variants with previously described hSNX16-associated markers Rab5 (early endosomes), Rab7 (late endosomes) and Rab11 (recycling endosomes) at the cell body, axon, and NMJ (**Fig. 8 A and Fig. S4 A-C**). In addition to the previously reported localization of dSNX16 at the NMJ to Rab5-positive endosomes (**Fig. 8 C and S4 A**; (Rodal et al., 2011)), dSNX16 localized at the cell body to both Rab5-positive and Rab7-positive puncta, suggesting different maturation states of dSNX16 containing endosomes between the nerve terminal and the cell body. dSNX16 did not co-localize with Rab11 in any region of the neuron, although strong diffuse Rab11 signal at the NMJ produced an artificially high PCC (**Fig. S4 B**). At the cell body, compared to wild type dSNX16, dSNX16^3A^ exhibited increased colocalization with Rab5-positive endosomes (PCC dSNX16 0.56 ± 0.09; dSNX16^3A^ 0.61 ± 0.10) and to an even greater extent with Rab7-positive endosomes (PCC dSNX16 0.62 ± 0.11; dSNX16^3A^ 0.77 ± 0.06). These CC-dependent changes in SNX16 association with Rab7-positive endosomes were not observed at the NMJ, where both SNX16 and SNX16^3A^ similarly reside on Rab5-positive endosomes (**Fig. 8, B and C**). To test whether the increased colocalization between dSNX16^3A^ and Rab5 and Rab7 at the cell body resulted simply from increased dSNX16^3A^ intensity, we quantified the intensity fraction of dSNX16 and dSNX16^3A^ in Rab5-positive (dSNX16 0.11 ± 0.05; dSNX16^3A^ 0.15 ± 0.06) or Rab7-positive compartments (dSNX16 0.09 ± 0.04; dSNX16^3A^ 0.18 ± 0.06) and observed comparable effects to our PCC measurements (**Fig. 8 D**). Finally, we did not find obvious colocalization between dSNX16 and our CC variants with Rab5, Rab7, or Rab11 in the proximal axon (**Fig. S4 C**), suggesting that transport carriers are rare among axonal dSNX16-positive structures, that their identity changes during transport, or alternatively that that cell body and NMJ SNX16 endosomes are independently derived from local membranes. These results indicate that in addition to altering endosome structure, abundance, distribution, and cargo sorting, dSNX16^3A^ also shifts the localization of SNX16 towards Rab7-positive late endosomes specifically at the cell body.

**Figure 8.**
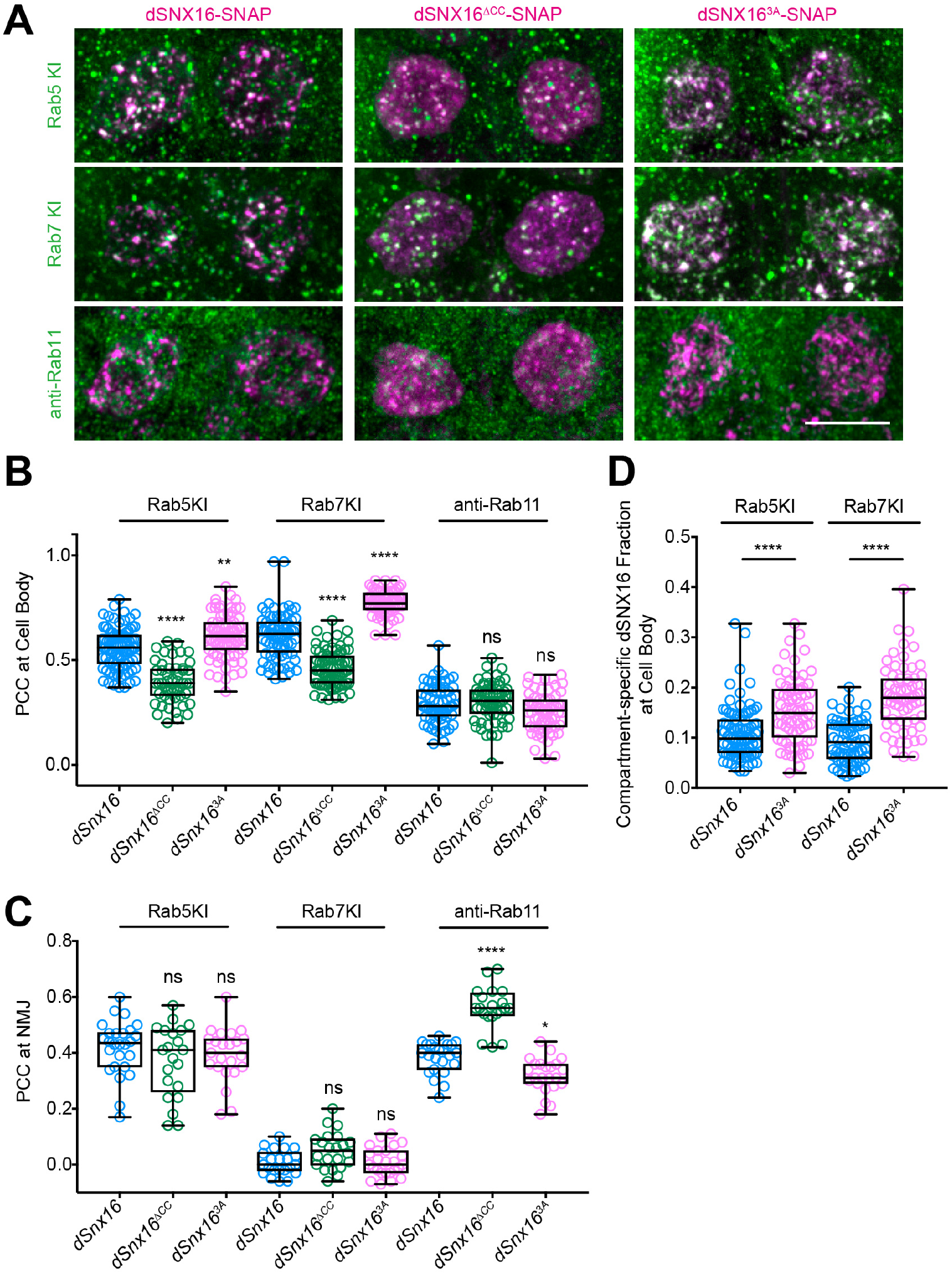
The dSNX16 coiled-coil domain regulates its localization to Rab7-positive endosomes at the cell body. dSNX16 CC variants localize primarily to Rab5-positive compartments at the NMJ, and at the cell body to both Rab5- and Rab7-positive compartments. dSNX16^3A^ showed increased colocalization with Rab5 and Rab7 specifically at the cell body. (A) Representative images of animals expressing VGlut-GAL4 driving the indicated dSNX16-SNAP^J^F549 transgenes. (B) PCC between dSNX16 and endogenous GFP-Rab5, YFP-Rab7, or anti-Rab11 immunoreactivity in the cell body. (C) PCC between dSNX16 and Rab5, Rab7, or anti-Rab11 at the NMJ. PCC for dSNX16^∆CC^ likely reflect its increased cytoplasmic localization. (D) dSNX16-SNAP^JF549^ intensity fraction in Rab5- and Rab7- positive compartments. Quantification is from at least 19 NMJs or 52 cell bodies, analyzed using Kruskal-Wallis test followed by Dunn’s multiple comparisons test. All images show 2D maximum intensity projections of confocal stacks. Data are shown as box-and-whisker plots with all data points superimposed. *p < 0.05, **p < 0.01, ****p < 0.0001. Scale bar, 10 μm.

### Rab5 activates dSNX16 endosome association, distribution, and remodeling in vivo

Our previous observation that dSNX16^3A^-expressing animals exhibit synaptic overgrowth, opposite from the effects of *dSnx16* null mutants, suggest that dSNX16^3A^ is a dominant active mutant (Rodal et al., 2011). Further our *in vitro* data indicates that SNX16^3A^ potentiates the normal oligomerization and membrane tubulation activity of SNX16, and does not behave as a neomorphic gain-of function. Therefore, we hypothesized that there may be mechanisms *in vivo* to activate dSNX16 towards a more dSNX16^3A^-like behavior for endosome association and deformation. We tested whether manipulations of Rab5 or Rab7 could activate dSNX16 based on our colocalization results with these GTPases. Strikingly, similar to dSNX16^3A^-expressing animals, overexpressing a constitutively active Rab5 transgene (Rab5^CA^) enhanced wild type dSNX16-GFP punctate localization at the proximal axon (where this phenotype is highly robust (**Fig. S3 D**)), and concomitantly enriched dSNX16 at the cell body (**Fig. 9 A-C**). By contrast, in dominant negative Rab5 (Rab5^DN^)-expressing animals dSNX16 exhibited primarily cytoplasmic localization (though not significantly different from the control by CoV), which is the opposite of dSNX16^3A^ and Rab5^CA^-expressing animals (**Fig. 9 B and Fig. S5 A**). dSNX16 levels were also elevated in all compartments in Rab5^DN^-expressing animals, perhaps due to its increased cytosolic distribution.

**Figure 9.**
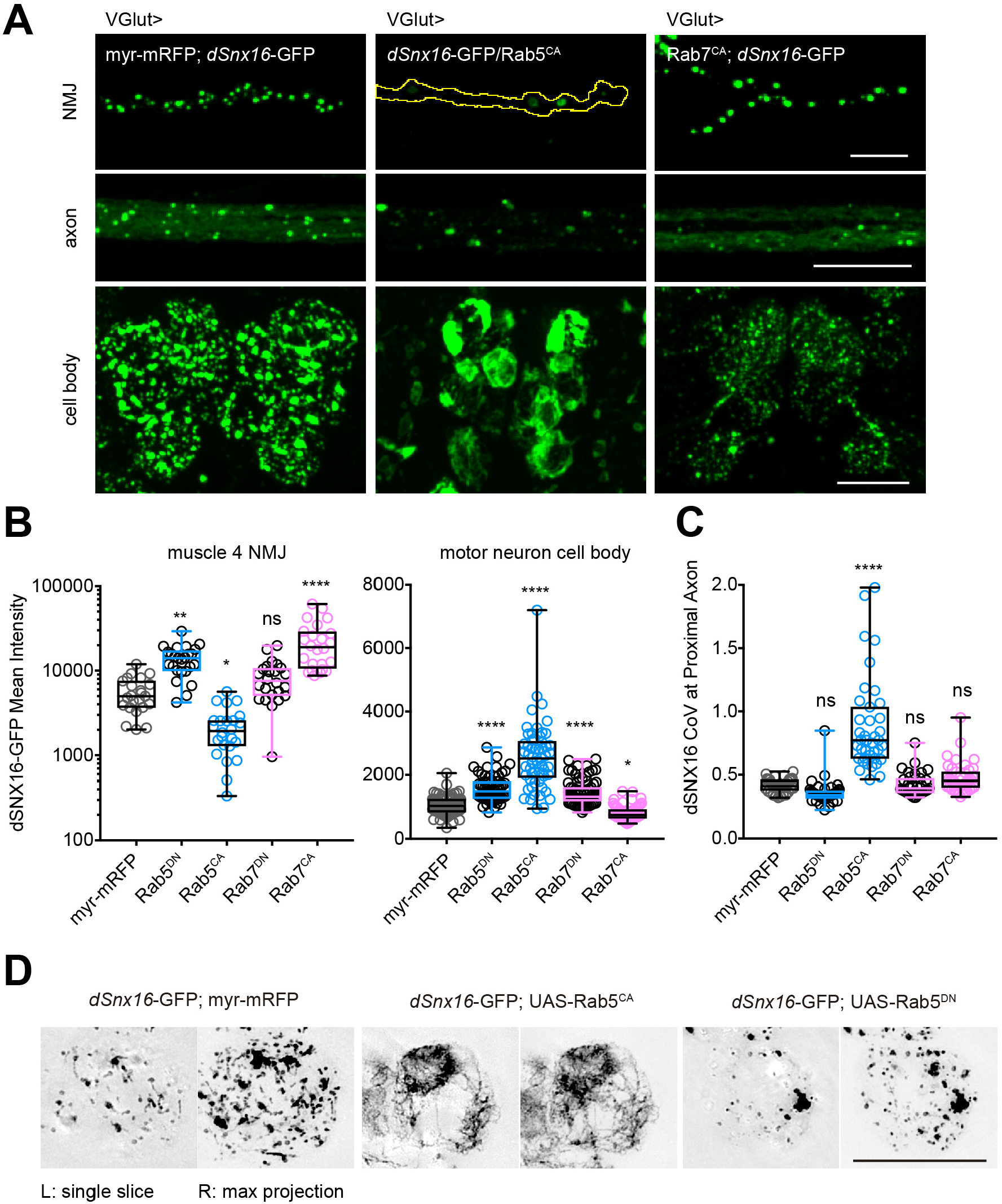
Rab GTPases regulate wild type dSNX16 endosome structure, localization and distribution in the neuron. (A) Representative images of animals expressing VGlut-GAL4-driven UAS-dSNX16-GFP together with Rab5^CA^, or Rab7^CA^ at the NMJ, the proximal axon, and the cell body. myr-mRFP serves as a UAS control. (B) dSNX16-GFP mean intensity quantification at the NMJ and the cell body. (C) Coefficient of variance (CoV) of dSNX16-GFP at proximal axon. (D) Representative SIM images of animals expressing VGlut-GAL4-driven UAS-dSNX16-GFP together with Rab5^DN^, or Rab5^CA^ at the cell body. Quantification is from at least 19 NMJs, 35 axons, or 52 cell bodies, analyzed using Kruskal-Wallis test followed by Dunn’s multiple comparisons test. All images show 2D maximum intensity projections of confocal stacks unless noted otherwise. Data are shown as box-and-whisker plots with all data points superimposed. *p < 0.05, **p < 0.01, ****p < 0.0001. Scale bar, 10 μm.

Next, since dSNX16 in Rab5^CA^-expressing animals exhibited dSNX16^3A^-like behaviors, we tested whether Rab5 can also modulate dSNX16 tubule-localization at the cell body (**Fig. 9 D**). Indeed, Rab5^DN^ and Rab5^CA^ dramatically affected dSNX16-decorated endosome structures. dSNX16 localized to exaggerated elongated threads in Rab5^CA^-expressing animals, consistent with more tubulated structures observed in dSNX16^3A^-expressing animals; while in Rab5^DN^-expressing animals, fewer tubulated dSNX16 endosomes were observed, similar to dSNX16^∆CC^.

Finally, we tested whether Rab7 similarly affected SNX16 endosome association and distribution. We found that dSNX16 was strongly concentrated at the NMJ in Rab7^CA^-expressing animals, consistent with previous findings (Akbergenova and Littleton, 2017), and was also largely depleted from the cell body (**Fig. 9, A and B**). By contrast, SNX16 was slightly enriched at the cell body in Rab7^DN^-expressing animals. Thus, Rab7 manipulations shifted the distribution of dSNX16 between NMJ and the cell body in the opposite direction of Rab5 manipulations, but did not affect dSNX16 endosome association, suggesting that Rab7-dependent endosome distribution effects occur downstream of SNX16 endosome association.

## Discussion

Here we propose a working model for SNX16 in regulating neuronal growth factor signaling by controlling endosomal tubulation and distribution (**Fig. 10**). At the molecular level, SNX16, via its CC domain, oligomerizes into higher-order assemblies that promote tubulation of PI(3)P-containing membranes. At the cellular level, the SNX16 CC domain is required for its endosomal localization in both mammalian cells and in *Drosophila* motor neurons. In these neurons, activation of Rab5 and CC-dependent self-association of SNX16 promote accumulation of SNX16 endosomes in Rab5- and Rab7-positive tubulated compartments at the cell body with the synaptic growth-promoting BMP receptor Tkv. Our results suggest that higher-order assembly of SNX16 on endosomes can control compartment identity and distribution to regulate the signaling activities of receptors in neurons.

**Figure 10.**
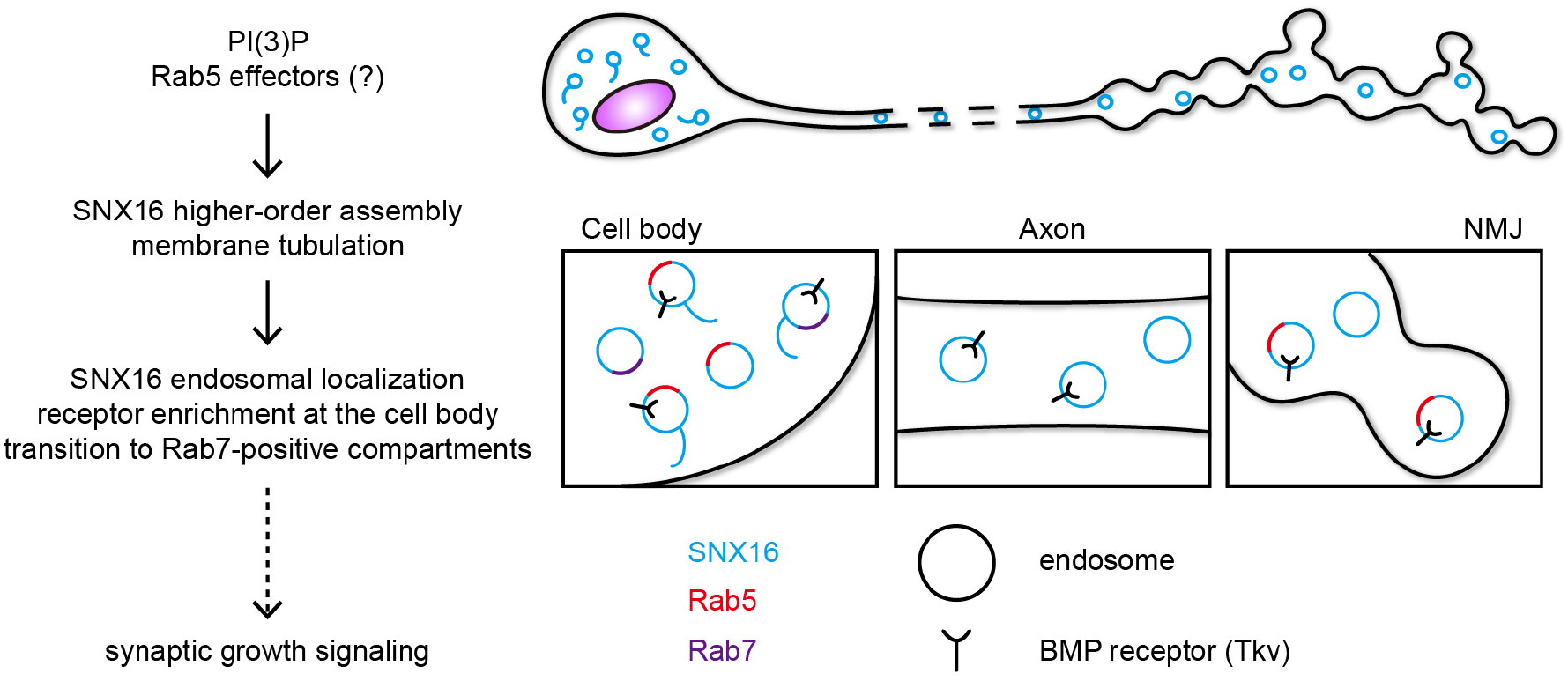
Working model for SNX16 function. The SNX16 CC domain controls its higher-order assembly and membrane tubulation capabilities. In *Drosophila* motor neurons, SNX16 oligomerization promotes its endosomal association, and redistribution from Rab5-positive endosomes at the NMJ to tubulated Rab5- and Rab7-positive endosomes at the cell body, containing the BMP receptor Tkv. SNX16 oligomerization may be regulated by Rab GTPases and/or PI(3)P levels *in vivo*.

### Membrane tubulation activity of SNX16

SNX16 is selectively localized to dynamic tubular late endosome structures *in vivo* (Brankatschk et al., 2011), but its function on these structures has been unclear. Here we provide direct evidence that SNX16 can generate membrane tubules via CC domain-mediated higher-order assembly (**Fig. 4**). Interestingly, SNX16 does not contain predicted membrane remodeling domains such as the amphipathic helices or BAR (Bin/Amphiphysin/Rvs167) domains found in the SNX-BAR subfamily (Pylypenko et al., 2007; van Weering et al., 2012), and therefore may deform membranes by one of several potential (and non-exclusive) mechanisms (McMahon and Boucrot, 2015; McMahon and Gallop, 2005). First, higher order assemblies of SNX16 may form a scaffold that imposes membrane curvature, as has been proposed for oligomers composed of Retromer and the PX-only protein SNX3 (Lucas et al., 2016). This model is supported by our cross-linking results and the reduced FRAP recovery we observe for SNX16 under conditions in which tubules form (**Fig. 2 and Fig. 3**), as well as previous results indicating multivalent self-association of SNX16 via PX and CC domains (Choi et al., 2004b). High-resolution EM studies of SNX16-decorated tubules will be required to resolve the oligomeric state and organization of these SNX16 assemblies. Second, the predicted unstructured regions in the N and C-termini of SNX16 (**Fig. 1 A**) may facilitate the scaffolding-induced membrane tubulation via protein crowding (Stachowiak et al., 2010; Stachowiak et al., 2012; Zeno et al., 2018). Third, the hSNX16 structure indicates that the hydrophobic residue F220, which is required for membrane binding, is positioned to penetrate into the membrane (Xu et al., 2017), potentially leading to wedging-induced membrane curvature. Indeed, our observation that higher-order assemblies of SNX16 are salt-insensitive for membrane binding suggest that these interactions may be mediated by such hydrophobic interactions (**Fig. 1 E and F**). Further biophysical studies of SNX16 membrane insertion will be required to test these hypotheses.

### Role of dSNX16 tubules in vivo

Consistent with its *in vitro* tubulation capability (**Fig. 4**) and the tubular localization of hSNX16 in HeLa cells (Brankatschk et al., 2011), we observed wild type dSNX16 on tubular structures that exclude the Tkv receptor at *Drosophila* motor neuron cell bodies (**Fig. 7 B**). The exclusion of Tkv from SNX16-decorated tubules stands in contrast to SNX-PXBAR decorated tubules, which colocalize with and sort cargo directly (Cullen and Steinberg, 2018), suggesting that different Sorting Nexins may either promote or restrict cargo transport in the endosomal pathway. The tubular localization of SNX16 is CC domain-dependent, and we observed increased numbers of tubular compartments in our hyper-oligomerized SNX16^3A^ mutant (**Fig. 6 B**). This mutant also promotes BMP receptor accumulation in endosomes (**Fig. 7**) as well as transcription of BMP target genes such as *trio* (Rodal et al., 2011). Taken together, our results raise the intriguing possibility that SNX16 tubules promote signaling by antagonizing BMP receptor downregulation, perhaps by removing other cargoes that would otherwise promote endosome maturation or lysosome fusion. Consistent with this hypothesis, overexpression of SNX16 blocks escape of viral RNA from MVBs (Le Blanc et al., 2005). Further, MVBs accumulate at SNX16^3A-^expressing NMJs (Rodal et al., 2011), and SNX16 is found on intermediate Rab7- and LAMP1-positive but not later LBPA- or CD63-positive endosomes (Brankatschk et al., 2011), suggesting that it may prevent endosome maturation.

### Role of dSNX16 in endosome distribution

In addition to regulating the structure of neuronal endosomes, we made the unexpected finding that self-association promotes SNX16 redistribution from Rab5-positive endosomes at the axon terminal to Rab5- and Rab7-positive endosomes at the cell body. One possibility is that SNX16-positive structures are independently generated at the cell body and the NMJ, similar to local recycling of Trk receptors (Ascano et al., 2009). Alternatively, SNX16 may play a role in long-range transport of endosomes from the NMJ to the cell body. In favor of this model, mammalian SNX16 co-purifies with retrogradely transporting, maturing compartments in neurons (Debaisieux et al., 2016); our results suggest that it may play an active role in this retrograde transport. Further, exit of Rab7-positive endosomes from the NMJ promotes BMP signaling (Liao et al., 2018), similar to TrkA receptor retrograde transport (Ye et al., 2018). Though we did not observe strong colocalization between dSNX16/dSNX16^3A^ and Rab5 or Rab7 in axons, it is possible that a small number of transport carriers are sufficient to mediate this redistribution. Taken together, these findings suggest that the endosome maturation/tubulation that occurs with SNX16 self-association is coupled with enhanced retrograde transport.

### In vivo regulation of dSNX16 oligomerization

dSNX16^3A^ exhibits an opposite synaptic growth phenotype compared to *dSnx16* null mutants (Rodal et al., 2011), and potentiates the normal *in vitro* activities of SNX16, suggesting that it functions as a dominant active mutant, and that *in vivo* mechanisms of wild type SNX16 regulation might similarly promote its oligomerization and endosome tubulation activities. Manipulation of Rab5 and Rab7 have previously been shown to affect dSNX16 levels at the NMJ (Akbergenova and Littleton, 2017), suggesting that these endosome regulators may control SNX16 activity. We found that GTP-locked Rab5^CA^ promotes endosome association, tubulation, and redistribution of wild type dSNX16 from the NMJ to the cell body (identically to dSNX16^3A^), while GDP-locked Rab5^DN^ produces the opposite phenotypes, suggesting that dSNX16 oligomerization and endosome tubulation may be regulated by Rab5. This may occur via direct effects on SNX16 CC interactions via an asyet-unknown Rab5 effector. Alternatively, Rab5 may regulate SNX16 activity by increasing PI(3)P levels (Shin et al., 2005), which can promote SNX16 membrane binding and remodeling *in vitro*. Separating these mechanisms will require developing tools to distinguish SNX16 CC-mediated oligomerization from PI(3)P binding. By contrast, manipulation of Rab7 did not directly phenocopy the effects of SNX16 oligomerization, and instead primarily resulted in redistribution of SNX16 between the cell body and the NMJ. Thus, Rab7 regulates transport of SNX16 endosomes but not assembly of SNX16 on these endosomes. Our previous results indicating that SNX16 acts downstream of the membrane-remodeling F-BAR/SH3 protein Nwk, which constrains synaptic growth signaling (Rodal et al., 2011), suggest that Nwk acts antagonistically to the Rab5 pathway to limit SNX16 activity. Further studies will be required to determine where Nwk and other potential SNX16-interacting factors act in this pathway, and to integrate opposing SNX16 regulatory mechanisms that control endosome maturation and distribution (**Fig. 10**).

## Materials and Methods

### Plasmid construction

Full-length human SNX16 (hSNX16) was amplified from Addgene plasmid #23617 and cloned into pcDNA3.1 using Gateway technology (Invitrogen). Mutations of hSNX16 were generated in this construct by site-directed mutagenesis. For expression in rat hippocampal neurons, hSNX16 and mutants were cloned into pCMV-myc. For expression and purification of hSNX16 variants, wild type and mutant hSNX16 were amplified via PCR and cloned into pGEX-6P (GE Healthcare) and pTrcHis-Xpress-SNAP vectors (Kelley et al., 2015). All constructs were verified by sequencing. pGEX-4T-hSNX1 was a gift from P. Cullen.

### Protein purification

BL21 DE3 cells transformed with the indicated constructs were grown to log phase and induced with 0.4 M IPTG at 18 °C for 12 hours. GST fusion proteins and His-Xpress-SNAP-tagged proteins were purified as previously described (Becalska et al., 2013; Kelley et al., 2015).

GST fusion proteins were purified from E. coli extracts on glutathione agarose resins (Thermo Scientific), followed by PreScission protease (GE Healthcare) cleavage at 4 °C for 16 hours. Released SNX16 variants were then gel filtrated through a Superdex 75 column in 20 mM Tris, 50 mM KCl, 0.1 mM EDTA, and 0.5 mM DTT, pH 7.5. Peak fractions were determined from SDS-PAGE analysis, concentrated, aliquoted, flash frozen in liquid N2, and stored at −80 °C. For BS3 crosslinking assays, proteins were gel filtered in 20 mM HEPES buffer instead of 20 mM Tris buffer.

His-Xpress-SNAP-tagged proteins were purified from E. coli extracts on nickel-nitrilotriacetic acid agarose beads (Qiagen), followed by gel filtration on a Superose 12 column (GE Healthcare) in 20 mM Tris, 50 mM KCl, 0.1 mM EDTA, and 0.5 mM DTT, pH 7.5. The concentrated peak fractions were supplemented with 1 mM DTT and incubated at 25 °C for 2 hours with a 5-molar excess of SNAP-Surface Alexa 488 (New England Biolab, Ipswich, MA). The final protein product was exchanged into 20 mM Tris, 50 mM KCl, 0.1 mM EDTA, and 0.5 mM DTT, pH 7.5 using a PD-10 desalting column (GE Healthcare), aliquoted, flash frozen in liquid N2, and stored at −80 °C.

### In vitro *liposome co-sedimentation and BS^3^ crosslinking assays*

Lipid co-sedimentation assays were conducted as previously described (Becalska et al., 2013). DOPS, PI(3)P, and Rhod PE were obtained from Avanti Polar Lipids (Alabaster, AL); DOPC and DOPE were obtained from Echelon Biosciences (Salt Lake City, UT). To generate liposomes, lipids were mixed in the indicated ratios and dried under a stream of argon gas, followed by 1 hour under vacuum. Lipid films were then hydrated in 20 mM HEPES pH 7.5, 100 mM NaCl for 2 hours at 37 °C to a final concentration of 1.5 mM.

In the co-sedimentation assays (all performed in triplicate), 10 μl of 1.5 mM liposomes (0.5 mM final concentration) with the indicated PI(3)P composition (0.05%, 1%, 2.5%, 5%, and 10%) were incubated with 20 μl of 3 μM protein (2 μM final concentration, in 20 mM HEPES pH 7.5, 100 mM NaCl) for 30 min at room temperature, followed by centrifugation at 18,000 × g for 20 min at 4°C. For salt sensitivity experiments, proteins and liposomes were incubated in 100mM NaCl or 400 mM NaCl for 45 min, or 100 mM NaCl for the first 30 min then in 400 mM NaCl for the following 15 min. Proteins were prespun under the same conditions before incubation with lipids. To measure membrane-bound fraction, pellets and supernatants were separated and equal amounts fractionated by SDS–PAGE, stained with Coomassie, and quantified by densitometry on a LICOR Odyssey instrument. Membrane-bound hSNX16 fraction was quantified as hSNX16 intensity in the pellet/ (hSNX16 intensity in the pellet + hSNX16 intensity in the supernatant)

For BS^3^ crosslinking assays, 10 μl of 1.5 mM 1% PI(3)P liposomes (0.5 mM final concentration) were incubated with 20 μl of 7.5 μM protein (5 μM final concentration) for 30 min at room temperature with BS^3^ (Thermo Fisher) at indicated concentrations (125 nM, 2.5 μM, 5 μM, 12.5 μM, and 25 μM in the liposome cosedimentation assay; 25 nM, 50 nM, 125 nM, 2.5 μM, 5 μM, 12.5 μM, and 25 μM in the HEPES buffer alone assay). 1 μl 1 M Tris was added to each reaction to stop crosslinking for 15 min at room temperature. Samples were then centrifuged, run on 7.5% SDS-PAGE gel, and imaged on LICOR Odyssey instrument as described in cosedimentation assays.

To account for the fact that in the BS^3^ crosslinking assays, hSNX16 pellet fraction and supernatant fraction were run on different gels, membrane-bound hSNX16 variants were quantified from the pellet and supernatant gels individually. In the pellet-fraction gels, hSNX16 intensity measured in no BS^3^ condition was considered the liposome-bound protein intensity. Intensity measurements were normalized to wild type hSNX16 intensity. In the supernatant-fraction gels, intensity measured in 0% PI(3)P liposome supernatant was considered the total protein intensity; intensity measured in no BS^3^ condition was considered the unbound protein intensity. Membrane-bound hSNX16 fraction was quantified as (total protein intensity – unbound protein intensity)/ total protein intensity. To control for background, pellet-fraction gels or supernatant-fraction gels of hSNX16 variants were stained in the same container and scanned in one image.

### In vitro *GUV assays and FRAP analysis*

GUVs were generated by electroformation in a Vesicle Prep Pro device (Nanion Technologies, Munich, Germany) or through gentle hydration in 5 mM HEPES and 300 mM sucrose, pH 7.5, as described (Kelley et al., 2015). Approximately 30 μM GUVs (300 nM PI(3)P) were mixed with 100 nM or 500 nM SNAP-tagged proteins in 10 mM HEPES, 150 mM KCl, pH7.5, incubated for 45min, and imaged in µ-Slide Angiogenesis devices (ibidi, Germany) on a Marianas spinning disk confocal system (see below).

To quantify the percentage of tubulated GUVs, the lipid channel (Rhod PE) was used to identify unilamellar vesicles. The protein channel (SNAP 488) was then used to count tubules for each GUV. The number of tubules reflected tubules observed in the whole Z-stack.

For FRAP experiments, 1% agarose in 10 mM HEPES 150 mM KCl, pH 7.5 was added to limit GUV mobility (0.33% final agarose percentage). GUVs were imaged for 6 min at 2-sec intervals, with a pause for bleaching after time point 10 (the 20th second, shown as time point 0 in graphs). Signal intensity over time from a non-bleached region of the same GUV was used to correct for photobleaching. Protein fluorescence was further normalized by subtracting the intensity at bleaching (time point 10) from all time points, and divided by prebleach intensity. Because lipid fluorescence recovered before the first time point was recorded (time point 10), it was not normalized, and was only corrected for photobleaching.

### Negative staining and electron microscopy

Liposomes for negative staining were generated at 1.5 mM as described in liposome co-sedimentation assays with 10% PI(3)P, and extruded through a 200 nm filter (Avanti Polar Lipids). Liposomes (0.5 mM) were incubated with protein for 30 min. Samples were applied to copper grids coated with continuous carbon, negatively stained with 2% uranyl acetate (JT Baker Chemical Co., Phillipsburg, NJ), and imaged using an FEI Morgagni 268 transmission electron microscope (FEI, Hillsboro, OR) operating at 80 kV and equipped with a 1k × 1k charge coupled device (CCD) camera (Gatan, Pleasanton, CA).

### Primary neuronal culture, transfection, and immunostaining

Dissociated rat hippocampal neurons were cultured on a feeder layer of astrocytes as previously described (Herzog et al., 2017). Briefly, a layer of confluent astrocytes was generated by plating the cells onto 12 mm glass coverslips coated with poly-D-lysine (20 μg/ml) and laminin (3.4 μg/ml) in 24 well plates. Dissociated hippocampi from E18 rat embryos were plated on this layer of astrocytes at a density of 80,000 cells/well, and grown in Neurobasal medium supplemented with B27 (Thermo Fisher) at 37 °C. Neurons were transfected at DIV 2 using the calcium phosphate transfection method (Xia et al., 1996) at 500 ng/well per plasmid, and fixed on DIV 7 with 4% PFA + 4% sucrose solution in PBS. pCMV-GFP plasmid was co-transfected with myc-hSNX16 variants to visualize individual neurons. Fixed neurons were then incubated with anti-Myc antibody (1:500) diluted in gelatin blocking buffer at 4 °C overnight in a humidified chamber, followed by 2 hours secondary antibody incubation at room temperature. Coverslips were then washed and mounted on glass microscope slides with Aquamount (Lerner Laboratories).

### Mammalian cell culture and transfection

Human adenocarcinoma HeLa and MCF7 cells, and human osteosarcoma U2OS cells (ATCC, Manassas VA) were grown in standard DMEM (Gibco, Life technologies, Grand Island NY) supplemented with 2 mM L-glutamine (Thermo Fisher Scientific) and 10% fetal bovine serum (FBS) at 37 °C with 5% CO2. Cells were seeded on collagen I-coated (50 μg/ml, Advanced BioMatrix, Carlsbad CA) 12 mm glass coverslips in 24-well plates before transfection. HeLa cells and U2OS cells were transfected at 40–50% confluence using polyethylenimine (PEI; Polysciences, Warrington PA) as previously described (Jansen et al., 2015). MCF7 cells were transfected at 70% confluence using Lipofectamine 3000 (Thermo Fisher Scientific) according to the manufacturer’s instructions. Cells were fixed in 4% PFA in PBS 24 hours after transfection. Coverslips were mounted on glass microscope slides with Aquamount (Lerner Laboratories).

### Fly stocks and generation of new strains

Flies were cultured using standard media and techniques. UAS-dSnx16-SNAP plasmids were constructed in pUAST-AttB (Brand and Perrimon, 1993) via Gibson assembly from previously generated UAS-dSnx16-GFP plasmids (Rodal et al., 2011). dSNX16 coil-coil mutations were generated by site-directed mutagenesis. Sequence-verified constructs were injected into w1118 flies at AttP40 (Markstein et al., 2008) at Rainbow Transgenic Flies (Camarillo, CA). Previously described fly stocks include VGlut-GAL4 (X) (Daniels et al., 2008), UAS-dSnx16-GFP and UAS-dSnx16^3A^-GFP (Rodal et al., 2011), UAS-Tkv-mCherry (Deshpande et al., 2016), GFP-Rab5KI (Fabrowski et al., 2013), YFP-myc-Rab7KI (Dunst et al., 2015), UAS-myr-mRFP (BL7118); UAS-Rab5^DN^ (Rab5^S43N/^; BL42703); UAS-Rab5^CA^ (Rab5^Q88L^; BL43335); UAS-Rab7^DN^ (Rab7^T22N^; (Assaker et al., 2010); UAS-Rab7^CA^ (Rab7^Q67L^; BL42707); UAS-Rab11^DN^ (Rab11^N124I^; (Satoh et al., 2005), and UAS-Rab11^CA^ (Rab11^Q70L^; (Emery et al., 2005).

### Imaging and statistical analysis

All GUV stacks were acquired at 0.5-μm steps using SlideBook software (3I, Denver, CO) on a Marianas spinning disk confocal system (3I, Denver, CO), which consists of a Zeiss Observer Z1 microscope (Carl Zeiss, Jena, Germany) equipped with a Yokagawa CSU-X1 spinning disk confocal head (Yokogawa, Tokyo, Japan) and a QuantEM 512SC EMCCD camera (Photometrics, Tucson, AZ) with a 100X (NA 1.45) oil immersion objective. Stack images were exported as tiffs into FIJI (National Institutes of Health, Bethesda, MD) for quantification as described in in vitro GUV assays and FRAP analysis.

To capture hSNX16 subcellular localization in fixed mammalian cells, spinning-disk confocal Z-stacks were collected at room temperature on an Andor spinning-disk confocal system, which consists of a Nikon Ni-E upright microscope equipped with a Yokogawa CSU-W1 spinning-disk head and an Andor iXon 897U EMCCD camera (Andor, Belfast, Northern Ireland). HeLa, U2OS, and MCF7 cells were acquired in NIS Elements AR software (Nikon, Melville, NY) with a 100X (NA 1.45) oil immersion objective at 0.2-μm steps. Cultured rat hippocampal neurons were acquired with a 60X (numerical aperture (NA) 1.4) oil immersion objective at 0.3 μm steps. To quantify coefficient of variance (CoV) of myc-hSNX16 intensity in the hippocampal neuron cell body, sum projections were generated from individual Z stacks with FIJI. Cell bodies were manually outlined in the GFP channel for transfected neurons. The mean fluorescence intensity and intensity standard deviation of myc-hSNX16 variants were then measured in that area to calculate CoV (standard deviation/ mean fluorescence intensity). To quantify hSNX16 and hSNX163A pixel intensity distributions in single-slice HeLa cells, cells were manually outlined. Images were normalized to the mean hSNX16 intensity in this area, and pixel intensity distributions were calculated using the Sixteen Bit Histogram plugin in Image-J. Data were binned as indicated and graphed using Prism software (GraphPad, La Jolla, CA).

For protein localization in Drosophila, flies were cultured at controlled density at 25°C. Wandering third-instar larvae were dissected in calcium-free HL3.1 saline (Feng et al., 2004) and then fixed in HL3.1 containing 4% formaldehyde before SNAP labeling or anti-Rab11 staining (clone 47 1:100 4 °C overnight; BD Biosciences). Larvae expressing dSNX16-SNAP variants were stained with 0.5 µM JF549-cpSNAP-tag ligand diluted in PBS for 2 min (Luke Laevis, Janelia Research Campus VA) (Kohl et al., 2014), followed by 1 hour α-HRP 647 staining (1:250; Jackson ImmunoResearch Laboratories, West Grove PA) at room temperature. For subcellular and colocalization analysis, type 1b NMJs from segment A2 and A3 on muscle 4 were imaged; 2 fields of proximal axons were imaged within 100 µm of the ventral ganglion; and 4 sets of MNISN-I cell bodies were imaged from the posterior end of the ventral ganglion (Choi et al., 2004a). Single NMJ stack images, single axon stack images, and single slice cell body images were manually cropped for intensity quantification.

Mean fluorescence intensity and CoV were performed on background-intensity-subtracted sum intensity projections at NMJ and proximal axon, and on single slices at the cell body, with the exception that CoV was quantified on max intensity projection at the proximal axon in **Fig. 6 B** because of the strong cytoplasmic localization. Pearson correlation coefficient (PCC) was calculated using Coloc2 (FIJI) on max intensity projections at NMJ and proximal axon, and on single slices at the cell body. To quantify compartment-specific dSNX16 fraction in **Fig. 8 C**, Rab5- or Rab7-positive compartments within the single slice cell body were outlined using ImageJ threshold setting Yen. dSNX16 intensity within that region was measured, and divided by the total cell body dSNX16 intensity.

For structured illumination microscopy (SIM) images at the cell body were collected at room temperature at 0.2-μm steps on a Nikon N-SIM instrument equipped with an Apo TIRF 100X (NA 1.4) oil immersion objective. Images were acquired using a violet-to-red diffraction grating at three angles and five phases of illumination, producing 15 raw images for SIM analysis, and reconstructed with default stack reconstruction setting in NIS Elements software.

All errors shown are mean ± SEM in XY and column graphs, as well as for quantifications reported in the text. All box-and-whisker plots are superimposed with individual data points. The box extends from the 25th to 75th percentiles with the median in the middle, and the whiskers mark the lowest data point to the highest. Data were tested for normality, and statistical significance was calculated in Prism software (GraphPad, La Jolla, CA) using one-way ANOVA followed by Tukey’s Test, Kruskal–Wallis tests followed by multiple Dunn comparisons (Kruskal–Wallis), or Mann-Whitney test as indicated in the figure legends. *p < 0.05, **p < 0.01, ***p < 0.001, ****p < 0.0001, ns not significantly different. Comparisons are to the leftmost genotype in each bar graph, unless indicated otherwise by horizontal bars.

## Acknowledgments

We are grateful to Peter Cullen and Luke Lavis for sharing reagents, the Bloomington Drosophila Stock Center (Indiana University, Bloomington, IN, NIH P40OD018537) for providing fly stocks; Steve Del Signore, Mugdha Deshpande, Suzanne Paradis, Tania Eskin, Silvia Jansen, and the Brandeis EM facility for technical assistance, Bruce Goode, M. Angeles Juanes, Mikael V. Garabedian, and Rodal lab members for helpful discussions. We thank Steve Del Signore, M. Angeles Juanes, Kate Koles, Biljana Ermanoska, and Cassandra Blanchette for comments on the manuscript. This work was supported NIH (R01 NS103967) and Pew Scholar awards to A.A.R. and by the Brandeis NSF MRSEC, Bioinspired Soft Materials (NSF-DMR 1420382).

The authors declare no competing financial interests.

## Author contributions

SW and AAR designed the study and experiments. SW conducted the experiments. SW and ZZ performed the analyses. SW and AAR wrote the manuscript.

